# Uncovering the Genomic and Phenotypic Landscape of Nitrogen-Fixing *Rhizobium miluonense* WD29 in Free-Living Conditions

**DOI:** 10.64898/2025.12.24.696422

**Authors:** Esaú De la Vega-Camarillo, Jossue Ortíz-Álvarez, Juan Alfredo Hernández-García, Lourdes Villa-Tanaca, César Hernández-Rodríguez

## Abstract

The endophytic bacterial strain *Rhizobium miluonense* WD29, isolated from the giant Mexican landrace maize Jala, demonstrates remarkable nitrogen fixation capabilities in free-living conditions. Through comprehensive genomic and phenotypic analyses, we characterized its unique attributes and potential applications in sustainable agriculture. Whole genome sequencing revealed a 6.8 Mb genome with 59.7% GC content, comprising 6,908 protein-coding genes, 3 rRNA genes, 46 tRNA sequences, and 146 insertion elements. Comparative genomic analysis showed that WD29 contains unique gene clusters associated with nitrogen fixation, biofilm formation, and plant growth promotion. The strain exhibits notable plant growth-promoting characteristics, including phosphate solubilization (26.1 ± 1.9 µg/mL), IAA production (19.7 ± 2.5 µg/mL), and multiple metallophore production capabilities (21.1-62.4% chelation for various metals). Significantly, *R. miluonense* WD29 demonstrates efficient nitrogen fixation in free-living conditions, with rates up to 21.7 ± 2.3 nmol h−^1^ of reduced acetylene at optimal galactose concentrations (1 g/L), coupled with substantial exopolysaccharide production (0.8 ± 0.076 g/mL). The strain’s genome harbors at least 9 genes involved in exopolysaccharide biosynthesis, likely contributing to its biofilm formation capability and enhanced nitrogen fixation efficiency. Additionally, WD29 shows remarkable environmental adaptability, possessing genes for heavy metal resistance and various stress responses. These findings highlight *R. miluonense* WD29’s potential as a valuable biofertilizer for sustainable agriculture, particularly for non-leguminous crops like maize, and demonstrate the importance of studying nitrogen-fixing bacteria isolated from traditional agricultural systems.

## 1. Introduction

Nitrogen is an essential element for plant growth, and most plants’ primary nitrogen source is derived from the soil [1]. However, nitrogen is often a limiting nutrient in the soil; plants usually compete with each other to acquire enough nitrogen for optimal growth [2-4]. Nitrogen-fixing bacteria help plants acquire nitrogen, converting atmospheric nitrogen into a form that plants can use [5-7].

The nitrogen cycle involves nitrogen fixation, nitrification, assimilation, and denitrification. *Rhizobium* species are nitrogen-fixing bacteria that form a symbiotic relationship with legume plants. Legumes’ root nodules convert atmospheric nitrogen (N_2_) into forms usable by plants, like ammonium (NH_4_^+^). This process enriches soil nitrogen, enhancing soil fertility and productivity [8,9].

*Rhizobium* species are nitrogen-fixing bacteria that form a symbiotic relationship with legume plants [10,11]. These bacteria are critical in nitrogen cycling and maintain soil fertility and productivity [11-13]. By understanding the genomics of *Rhizobium* species, we can better understand the genetic mechanisms underlying nitrogen fixation and other phenotypic traits, such as symbiotic compatibility with legumes and other plant hosts [15-17].

Nitrogen-fixing is a remarkable biological process that converts atmospheric gaseous nitrogen (N2) into an assimilable form that *Rhizobium*-plant symbiosis can utilize [18]. This symbiotic relationship between rhizobia and plants such as leguminous is crucial for sustainable agriculture and ecosystem functioning. *Rhizobium* species form nodules on the roots of these plants, providing a specialized microenvironment for nitrogen fixation. Inside the nodules, nitrogen-fixing is carried out by nitrogenase, which catalyzes the conversion of nitrogen gas into ammonia (NH3), a form of available inorganic nitrogen source [19]. This process benefits the host plant by providing a vital nutrient and contributes to the overall enrichment of soil fertility [1].

Nitrogen-fixing in *Rhizobium* exhibits distinct behaviors under symbiotic and free-living conditions. *Rhizobium* establishes a mutualistic relationship with leguminous plants in symbiotic conditions, forming nodules on their roots [18]. On the other hand, *Rhizobium* also exhibits a free-living lifestyle outside of symbiotic associations [20]. Although a few works have documented this phenomenon, in free-living conditions, *Rhizobium* can survive and persist independently in the soil or rhizosphere, contributing to the nitrogen cycle by mineralizing organic nitrogen compounds and releasing ammonium into the soil.

The genomes of most *Rhizobium* species contain *nif* gene operons responsible for biological nitrogen fixation, which has evolved over millions of years and is likely shaped by the coevolutionary history of *Rhizobium* species and their legume hosts [21-23]. By analyzing *nif* gene operons, it is possible to obtain more insights into the evolutionary processes that have shaped these symbiotic relationships and potentially use this knowledge to develop more sustainable agricultural practices [24].

*R. miluonense* is a *Rhizobium* species found in various soils worldwide, including Australia, Brazil, Canada, and China. It is known for its nitrogen-fixing ability, but relatively little is known about its genome and phenotypic traits [25-27].

This work comprehensively analyzes the genome and phenotypic traits of the strain *R. miluonense* WD29 isolated from Jala landrace maize rhizospheric soil in Nayarit, México. The aim was to understand better the phenotypic and genotypic characteristics of our strain of study associated with nitrogen fixation in free-living conditions and other phenotypic traits associated with plant-grow promotion. Specifically, we explored the genomic content related to nitrogen fixation, carbon metabolism, and stress response in the

*R. miluonense* WD29 genome project, besides its ability to tolerate various environmental conditions and form biofilms. We also confirm its ability to fix nitrogen in free-living conditions, with a rate of up to 21.7 nmol h-1. Our findings provide important insights into the potential applications of *R. miluonense* WD29 in agriculture and environmental science and highlight the importance of studying nitrogen-fixing bacteria for sustainable agriculture and ecosystem health.

## 2. Materials and Methods

### 2.1 Bacterial strain and growth conditions

The rhizospheric bacterium *R. miluonense* WD29 was previously isolated from the Jala landrace maize rhizosphere [28]. The strain was routinely cultured in R2A medium at 28 ± 1°C for 48 h with constant shaking (150 rpm). For long-term storage, bacterial cultures were maintained in 70% (v/v) glycerol at -70°C.

### 2.2. Genomic Analysis

#### 2.2.1. DNA Extraction

Genomic DNA extraction was performed following the CTAB protocol [29]. The concentration and desired purity relation 260/280 was determined in a Nanodrop. The integrity was analyzed in an agarose electrophoresis gel at 1%. The approved DNA sample for WGS sequencing was chosen according to the quality criteria of concentration, purity, and integrity requested by the sequencing company for complete genome sequencing.

#### 2.2.2. Whole Genome Sequencing and Assembly

The whole genome sequencing of R. miluonense WD29 was carried out using the Illumina HiSeq platform with Nextera XT Library Prep (Novogene, North America). Quality control of raw data was analyzed with FastQC v. 0.11.9 [30]. Raw reads were filtered and trimmed with Trimmomatic [31]. The de novo assembly was performed with SPAdes v. 3.13.1 using default parameters [32]. Assembly quality evaluation was conducted with QUAST v. 5.0.2 [33]. The Rhizobium group assessed Genome completeness and annotation with BUSCO version 5.4.6 [34]. Genome annotation was performed using the RAST server v.2.0, Prokka version 1.14.6, and the NCBI Prokaryotic Genome Annotation Pipeline [35-37]. Secondary metabolite clusters were identified using antiSMASH bacterial version v. 6.0 [38] and PRISM [39].

### 2.3. Phylogenomic and Comparative Analysis

The genome was analyzed using the Average Nucleotide Identity (ANI) algorithm, specifically ANIb [40]. Genome sequences were compared to NCBI RefSeq database entries using PyANI v0.2.8 and visualized using R’s pheatmap package [41]. Taxonomic assignment utilized the M1CR0B14L1Z3R platform with parameters: maximal e-value cutoff: 0.01, identity minimal percent cutoff: 90.0%, and minimal percentage for core: 100.0% [42].

The comparative genomic analysis included 38 strains from the genus *Rhizobium*/*Agrobacterium*. Genes involved in plant-microorganism interactions were analyzed, including those involved in phosphorus and zinc mobilization, heavy metal tolerance, phytohormones, biofilm formation, siderophore biosynthesis, biological control, quorum sensing and quenching, biological nitrogen fixation, nodulation factors, cell motility, cellulases, and antibiotic resistance. Data visualization was performed using TBtools-ll v1.108 [43].

Orthologous clustering analysis used Orthovenn3 with the Orthomcl algorithm [44]. Genome synteny and collinearity analysis employed MAUVE [45]. Gene gain/loss analysis was conducted using CAFE5, using the time of divergence between R. miluonense and Rhizobium tropici (50.85 Mya) [46].

### 2.4. Plant Growth-Promoting Characterization

Phosphate solubilization, ACC deaminase detection, indoleacetic acid production, and siderophore production were determined according to previously reported protocols [28]. Detection of extracellular enzymes (chitinases, cellulases, and proteases) followed established protocols [47].

### 2.5. Biological Nitrogen Fixation Assays

Nitrogenase activity was estimated using acetylene reduction assay with gas chromatography [48]. The bacterial strain was grown in sealed bottles containing 5 mL of semisolid BMGM medium with varying galactose concentrations (0.25, 0.5, 0.75, and 1 g/L). Bacterial suspensions were adjusted to 1×10^8^ CFU/mL, with *Klebsiella variicola* 6A3 as a positive control [28]. After 48h incubation at 28ºC, 400 µL of headspace gas was replaced with acetylene and incubated for 6h. Analysis was performed using a Perkin Elmer Clarus 580 gas chromatograph.

### 2.6. EPS Biosynthesis Pathway Analysis

KEGG database searches employed BlastKOALA and GhostKOALA tools [49,50]. Protein domain analysis used InterProScan 5 [51]. Gene function verification involved alignment with known EPS biosynthetic genes using Clustal Omega, visualized with Jalview [52,53].

Protein sequences were obtained from the genome sequence of *R. miluonense* WD29. To predict the 3D structures of the proteins encoded, we used AlphaFold2, a deep learning-based protein structure prediction software. To validate the accuracy of the Alphafold2 predictions, we compared them to experimentally determined structures of homologous proteins, where available. Homologous protein structures were obtained from the Protein Data Bank (PDB) using the BLASTP algorithm to search for proteins with significant sequence similarity [54,55].

### 2.7. Statistical Analysis

All experiments were performed in triplicate. Data are presented as mean ± standard deviation. Statistical significance was determined using one-way ANOVA followed by Duncan’s test (P ≤ 0.005). Statistical analyses were performed using R version 4.1.0.

## 3. Results

### 3.1 Genomic features

The genome of *R. miluonense* WD29 had a size of 6.8 Mb and is composed of 103 contigs with an N50 contig length of 108 kb, 3 rRNA genes, 46 tRNA sequences, and 146 insertion elements (IS). The GC content of the genome is 59.7%, similar to that of other *Rhizobium* species. A total of 6,908 protein-coding genes were predicted; 5,284 genes were assigned a COG functional category, with the most significant categories being amino acid transport and metabolism, transcription, and carbohydrate transport and metabolism (Figure 1).

**Figure 1.**
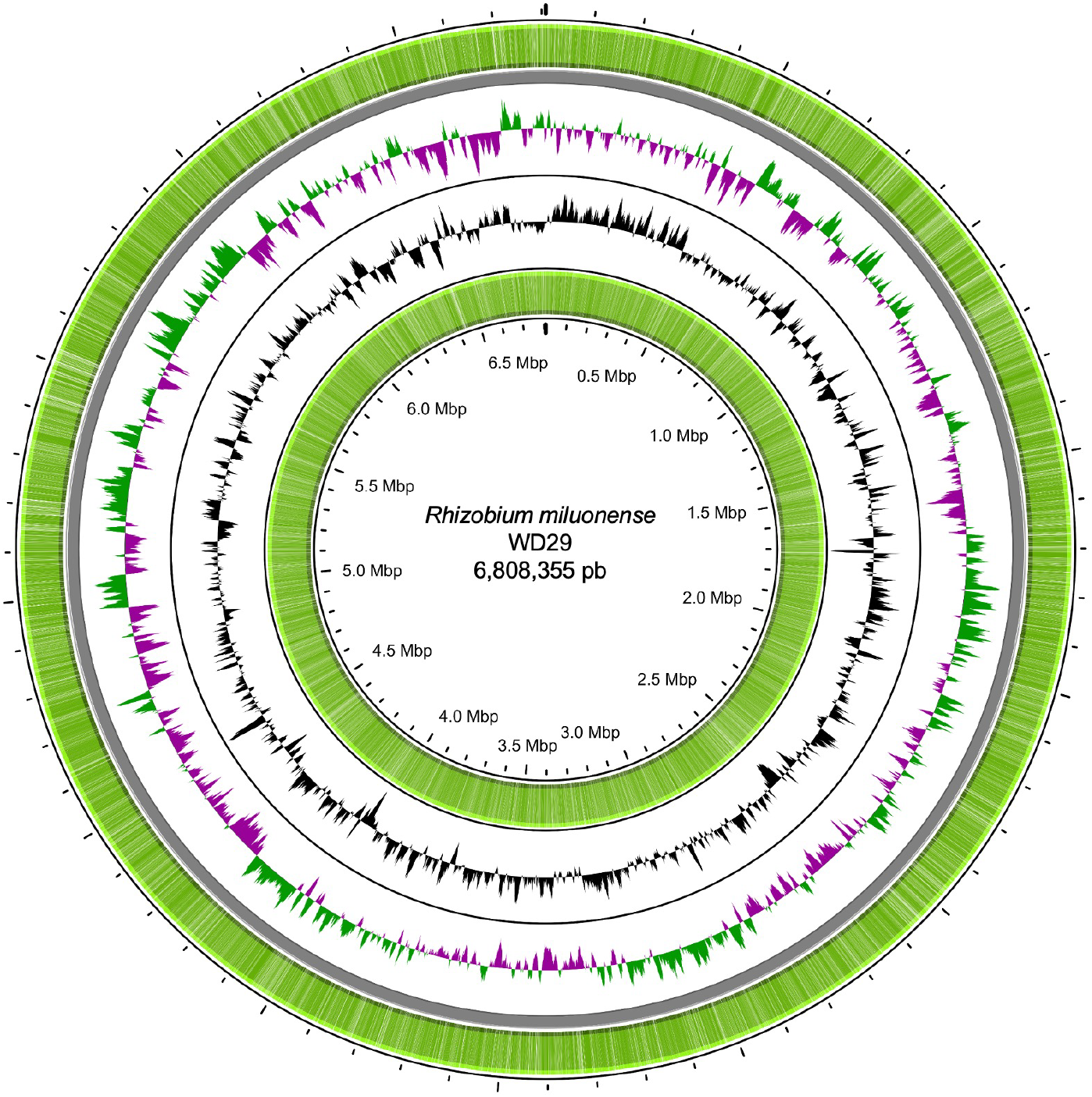
Circular representation of the *Rhizobium miluonense* WD29 genome. The outermost and innermost circles represent the coding sequences of the leader and complementary chains of the *R. miluonense* WD29 genome; the next circle corresponds to the positive GC bias in green and negative in purple, and the next circle represents the GC content.

### 3.2 Phylogenomic, comparative, and functional genomics

The core-genome-based phylogenetic tree clustered to the WD29 strain with the type strain of *R. miluonense* HAMBI 2971 (IMG-taxon 2617270742), confirming the identity of our strain of study as a strain belonging to *Rhizobium miluonense* species (Figure 2). Also, the phylogeny showed that *R. miluonense* WD29 is a member of the more prominent family of Rhizobiaceae, which also includes other nitrogen-fixing bacteria that form symbiotic relationships with legumes, such as *Rhizobium leguminosarum* and *Sinorhizobium meliloti* [56]. Within the family Rhizobiaceae, *R. miluonense* forms a clade with other strains of *Rhizobium*, including *Rhizobium freirei, Rhizobium multihospitium*, and *Rhizobium tropici*. This clade is supported by a high bootstrap value (100), which suggests that these strains are closely related.

**Figure 2.**
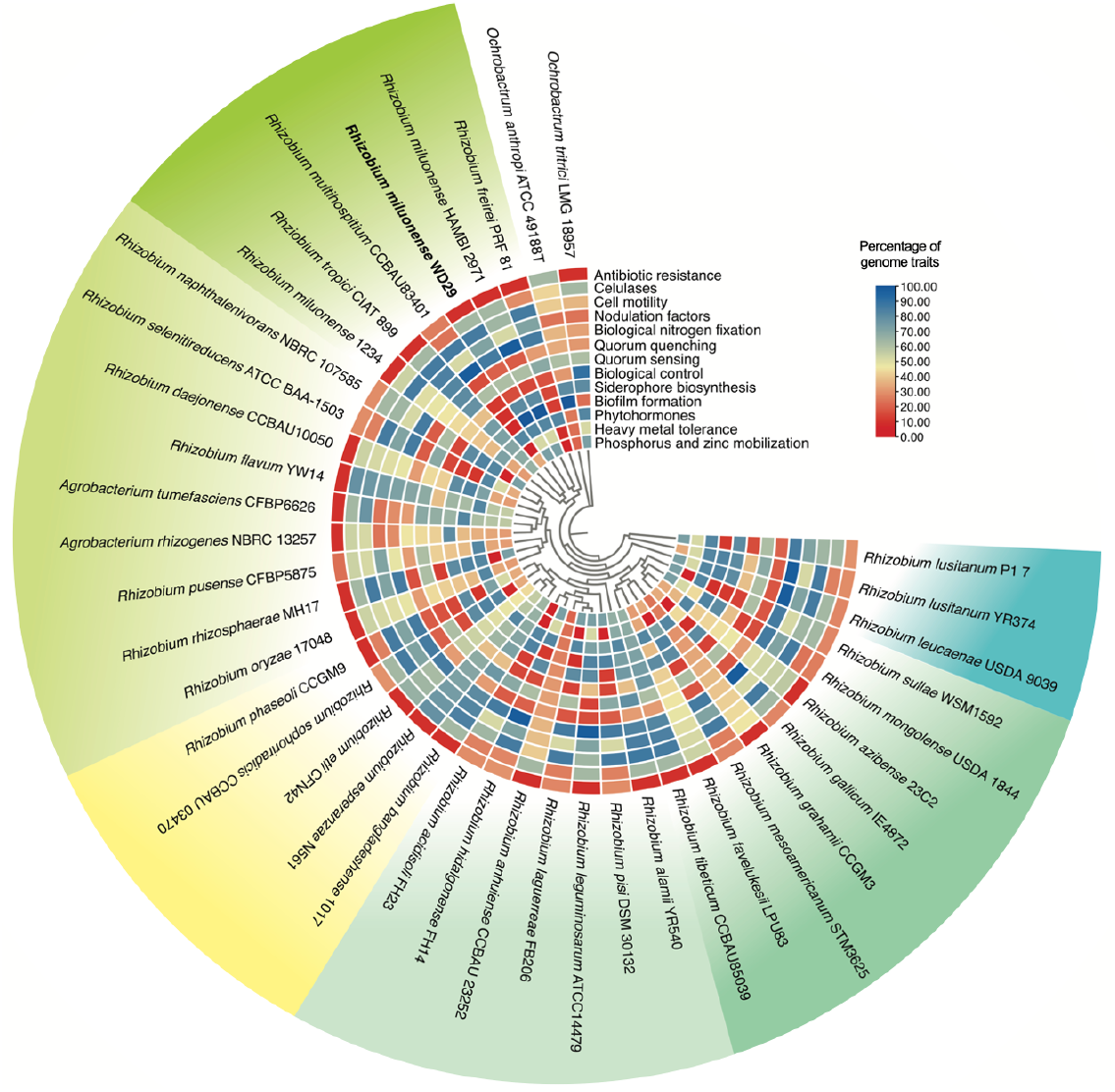
Phylogenomic and comparative genomic analysis between *R. miluonense* WD29 and the different species of the genus *Rhizobium*. The phylogenomic was constructed from the core-genome comparison using the maximum likelihood method, and the genomic content was expressed as a percentage of genes associated with a relevant trait in the plant-microorganism interaction. The color scale in the chart represents a gradient of gene abundance. Blue colors represent a greater abundance of genes, while red colors represent low abundance.

When the genomic content based on essential activities in the plant-microorganism interaction was analyzed, we denoted that, in general, the clade in which *R. miluonense* WD29 is located contains a higher content of genes associated with mobility and nitrogen fixation, especially *R. miluonense* species contain more genes involved in biofilm production and low content of genes involved in biological control compared to their close relatives. In general, the species of the Rhizobiaceae family contain a low genomic content associated with antibiotic resistance. In particular, the WD29 strain exhibited genes related to resistance to fluoroquinolone and tetracycline (Figure 2).

Analysis of orthologous clusters showed that *R. miluonense* WD29 and its closely related species share a core-genome shaped of 4011 clusters of orthologous groups (COGs) (Figure 3B). Both *R. miluonense* strains share 680 COGs, mainly associated with primary metabolism, secondary metabolism, biofilm formation, and nitrogen metabolism, which may be involved in the plant-host interaction. Interestingly, the WD29 strain contains 7 accessory COGs related to biological functions such as DNA integration and carbon source transport and 4 COGs with unknown functions. Also, WD29 and its most closely related species, *R. freirei*, share 4 CGOs involved in transcription, antibiotic resistance and two COGs with unknown functions, which are not present in *R. miluonense* type strain. Both *R. miluonense* strains share 431 COGs with *R. freirei*. Gene gain/loss analysis displayed that Both *R. miluonense* strains had different expansions and contraction rates (Figure 3A). For instance, the results show that the WD29 strain suffered more expansion events and fewer contractions (+9/-23) than the *R. miluonense* type strain (+2/-53) (Figure 3A). Nonetheless, both *R. miluonense* strains exhibit fewer events of gene gain and more gain losses than *R. freirei, R. tropici*, and *R. multihospitium* (Figure 3A). MAUVE analysis showed that both *R. miluonense* genomes maintain a very well-conserved synteny in almost the entire genome (Figure 3C), with few non-syntenic blocks displayed as white gaps at the terminal regions. On the other hand, the genome of *R. freirei* exhibits several rearrangements and inversions compared with *R. miluonense*.

**Figure 3.**
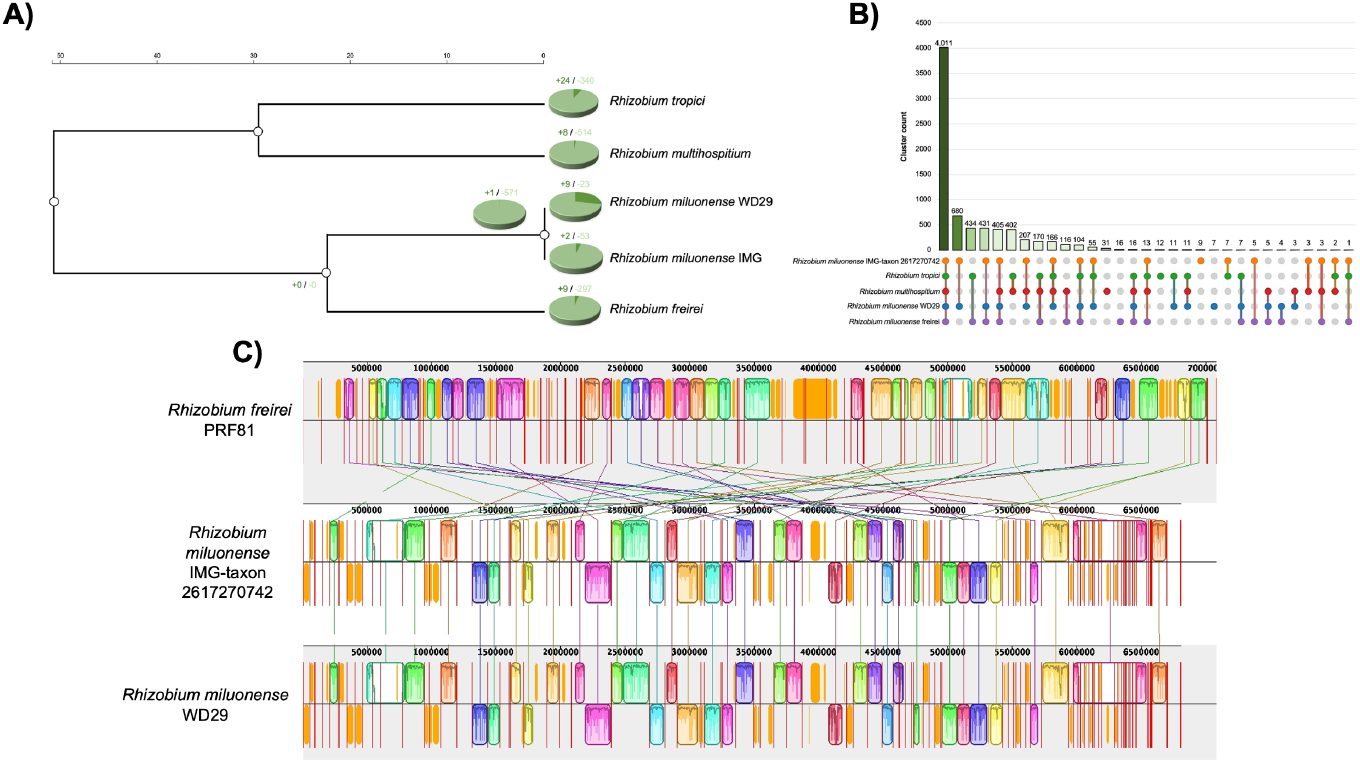
Comparative genomics among *R. miluonense* WD29 and its most closely related *Rhizobium* species. **A)** Gene gain/loss analysis among *R. miluonense* WD29 and its closely related *Rhizobium* species. Pie charts above each taxon represent the expansion and contraction occurring in each genome. The tree represents the phylogenetic relationship among all the taxa chosen for the analysis. The scale represents the estimated theoretical time of divergence computed for *Rhizobium* species. **B)** UpSetR comparison plot among Clusters of Orthologous Groups (COGs). The numbers above the vertical bars represent the abundance of shared COGs among the closely related *Rhizobium* species. Horizontal bars represent the overall content of predicted proteins in each genome. The dot plot represents a graphic representation of the binary presence/absence of COGs. **C)** Rainbow shades represent genomic blocks. Variable regions are described as white gaps. Connected lines represent the shared well-conserved regions among genomes.

### 3.3 Detection of putative Plant-Growth Promotion Compounds (PGPC)

Phenotypic analysis showed that *R. miluonense* WD29 is capable of producing some relevant compounds and enzymes cataloged as PGPC, such as phosphate solubilization, IAA production, metallophores for different metal ions, chitinases, cellulases, and proteases, which were quantified and the results are shown in Table 1.

**Table 1.**
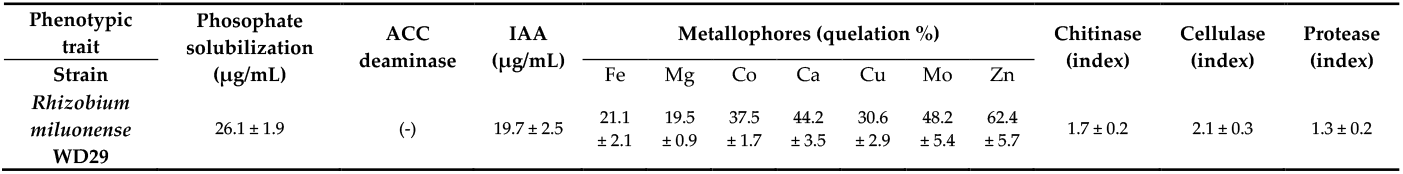
Relevant phenotypic traits associated with the plant-microorganism interaction of *R. miluonense* WD29.

### 3.4 Biological nitrogen fixation

The concentration of galactose (g/L), the diameter of colonies (mm), nitrogen fixation (nmol h-1), and exopolysaccharide (EPS) production (g/mL) were measured in this study (Figure 4). The galactose concentration increased from 0.25 g/L to 1 g/L, resulting in a corresponding increase in the diameter of colonies from >1 mm to 8.1 mm. The relationship between galactose concentration and colony size appeared to be concentration-dependent. Nitrogen fixation was evaluated, and the results showed that it was not detected (ND) at the lowest galactose concentration (0.25 g/L). However, as the galactose concentration increased to 0.5 g/L, nitrogen fixation was observed at 7.3 ± 0.8 nmol h-1. At higher galactose concentrations (0.75 g/L and 1 g/L), nitrogen fixation rates increased to 10.4 ± 1.8 nmol h-1 and 21.7 ± 2.3 nmol h-1, respectively. EPS production was also assessed, and it was found to be no detectable (ND) at the lowest galactose concentration (0.25 g/L). However, as the galactose concentration increased to 0.5 g/L, EPS production was measured at 0.1 ± 0.019 g/mL. Further increases in galactose concentration (0.75 g/L and 1 g/L) led to significant increases in EPS production, reaching 0.3 ± 0.023 g/mL and 0.8 ± 0.076 g/mL, respectively.

**Figure 4.**
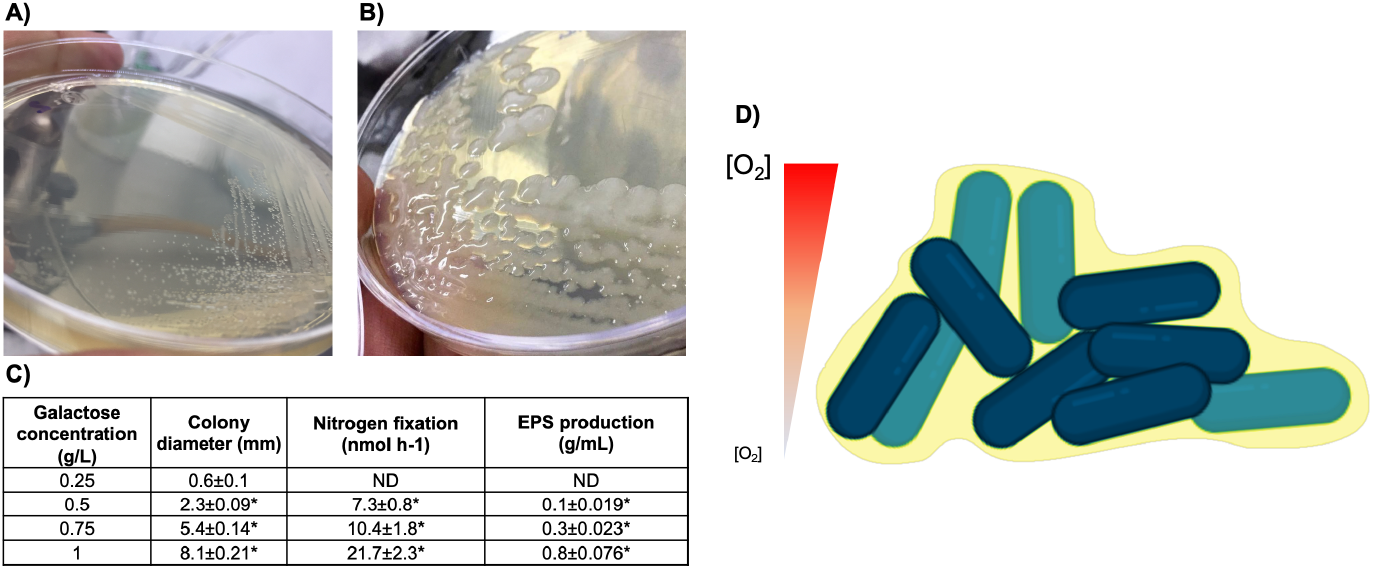
Biological nitrogen fixation in vitro of *R. miluonense* WD29. **A)** Colonial morphology of *R. miluonense* WD29 in LGI medium with 0.25 g/L galactose, **B)** Colonial morphology of *R. miluonense* WD29 in LGI medium with 1 g/L galactose, **C)** relationship of galactose concentration in the medium with colony diameter, EPS production, and nitrogen fixation rate; each assay was performed in triplicate, * means statistically significant difference compared to the control (0.25 g/L of galactose) with Duncan’s test (*P≤ 0.005), **D)** Representative scheme showing the mechanism of action of EPS in biological nitrogen fixation.

### 3.5 EPS biosynthesis

In the genome of *R. miluonense* WD29, we identified at least 9 genes involved in exopolysaccharide biosynthesis (Table 2), which were compared with other synthesis pathway genes in other members of the *Rhizobium* genus. By comparing the proteins engaged in previously reported EPS biosynthesis pathways, we can assume that this bacterium follows the biosynthesis pathway via exo genes, as reported for other species of *Rhizobium* and *Sinorhizobium* (Figure 5).

**Table 2.** Genes involved in EPS biosynthesis in *Rhizobium miluonense* WD29By comparing the proteins engaged in previously reported EPS biosynthesis pathways, we can assume that this bacterium follows the biosynthesis pathway via exo genes, as has been reported for other species of the genus *Rhizobium* and *Sinorhizobium*. (Figure 5).

| Gen | Protein | Genomic location |
| --- | --- | --- |
| *wbaP* | Undecaprenyl-phosphate galactosephosphotransferase | 52706-53398 |
| *wbpY* | Glycosyl transferase | 6794-8856 |
| *agpA* | ABC-type sugar transport system, periplasmic component | 103655-103753 |
| *ptsG* | ABC-type sugar transport system, permease component | 86150-86971 |
| *exoP* | Exopolysaccharide polymerization/transport protein | 89588-91972 |
| *exoZ* | Exopolysaccharide production protein | 73276-74283 |
| *exoQ* | Exopolysaccharide production protein | 74280-75572 |
| *exoF* | Exopolysaccharide production protein | 75582-76856 |
| *exoR* | Exopolysaccharide production negative regulator precursor | 171123-171890 |

**Figure 5.**
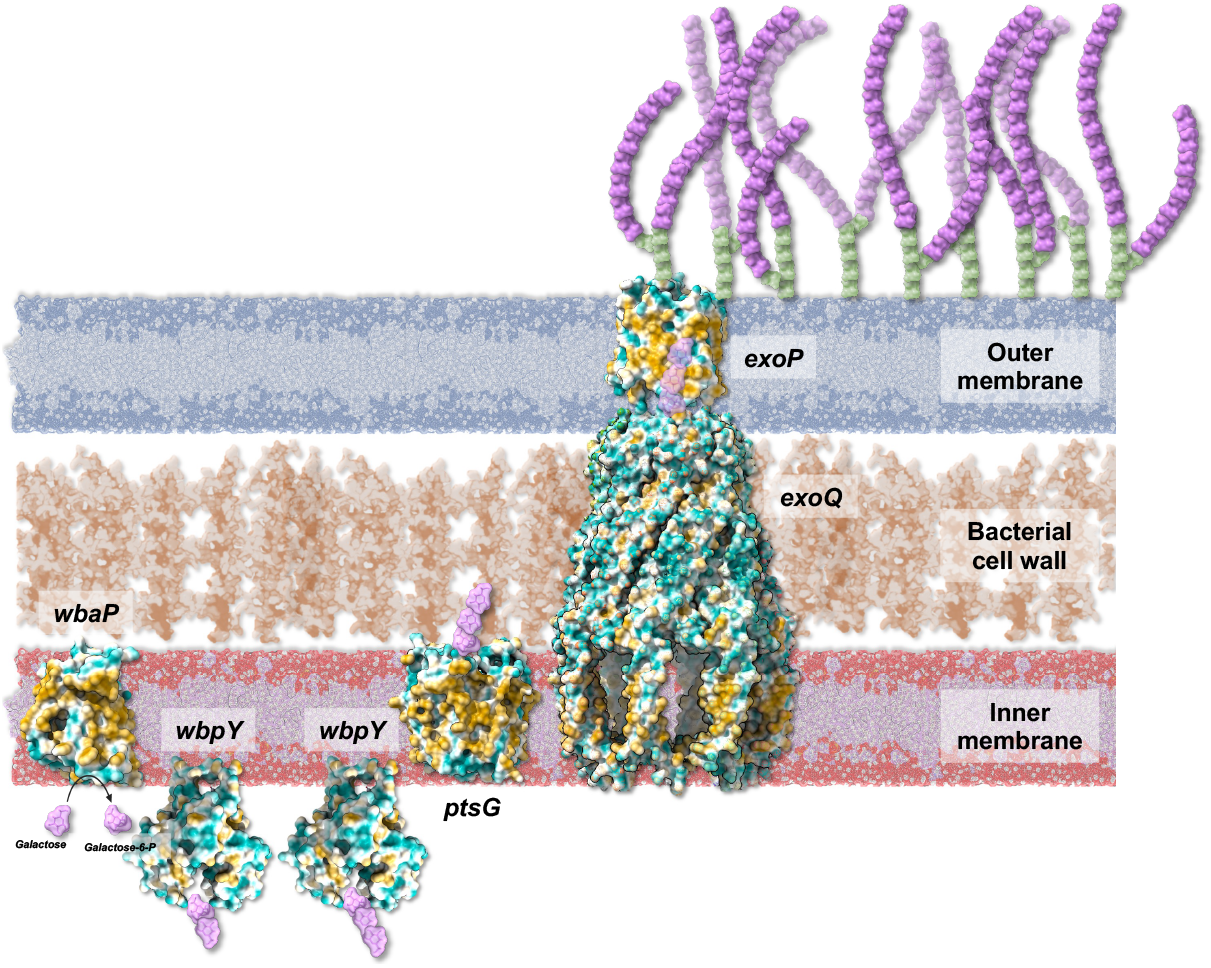
Representative scheme of LPS biosynthesis in *R. miluonense* WD29. Deduction of protein function was performed by homology comparison with previously described proteins for other members of the genus. The protein structure prediction was performed using AlphaFold2, and the visualization and realization of the scheme were performed in BioRender and ChimeraX.

## 4. Discussion

In this study, we explored the genome and phenotypic features associated with Plant Growth Promotion (PGP) of *R. miluonense* WD29 due to its peculiarity of free-living nitrogen fixation (FLNF), a characteristic that has not been explored in depth in this particular genus, since most of the research is based on nitrogen-fixing performed by the *Rhizobium* genus in symbiotic conditions within the nodules that form with legumes [18,57-59]. Previous studies report that *Rhizobium* sp. RC3200 possesses free-living nitrogen fixation forming specialized nongrowing cells, a common trait observed in Cyanobacteria [20]. However, this phenomenon’s molecular mechanisms and genomic features still need to be explored. This is the first study to investigate the free-living nitrogen fixation property using genomic methods in *Rhizobium* species.

Identifying 6,908 protein-coding genes in the genome of *R. miluonense* WD29 provides valuable information about its metabolic capabilities and potential functions. The genes were assigned to COG functional categories, with the most significant categories related to amino acid transport and metabolism (lysine, threonine, methionine, cysteine, aromatic amino acids, and derivatives), transcription (transcriptional regulators and repressors), and carbohydrate transport and metabolism (fermentation, oligosaccharides, and sugar alcohol). These results suggest that *R. miluonense* WD29 has diverse metabolic capabilities, allowing it to utilize various carbon and nitrogen sources. Identifying many genes related to amino acid transport and metabolism suggests that this bacterium is highly required for these nutrients and can use them efficiently. Nonetheless, carbon and nitrogen assimilation tests in further works are necessary to confirm this assertion.

The WD29 strain exhibited a higher content of genes associated with nodulation than the *R. miluonense* type strain. The nodulation factor genes allow Rhizobia to penetrate and infect the root, besides nodule formation [60]. Also, *R. miluonense* has demonstrated an efficient symbiotic performance in contact with *Phaseolus vulgaris* [61]. According to our results and previous in vivo findings, increasing nod factor genes in the WD29 strain could promote better nodulation than the *R. miluonense* type strain. We also observed that *R. miluonense* possesses more genes related to Plant Growth Promotion (PGP) activities than *R. freirei* and *R. multihospitium*.

Moreover, both *R. miluonense* strains exhibited considerable genes associated with Heavy Metal Resistance (HMR). Our species of study has been demonstrated to grow in the presence of chromium [61] and can solubilize hydroxyapatite for phosphorus bioconversion [63]. Previous reports support our findings and evidence a strong correlation between phenotype and genomic features.

Although the WD29 strain showed high ANI values compared to the reference genome of *R. miluonense*, we discovered significant differences in the genomic features. Both *R. miluonense* strains share 600 exclusive COGs; notably, WD29 displayed 7 unique COGs. The exclusive COGs of *R. miluonense* had functional assignments to primary metabolism, secondary metabolism, biofilm formation, and nitrogen metabolism that may be involved in the plant-host interaction. The WD29-specific COGs are associated with tetrahydrofolate conversion, DNA integration, carbon source transport, and unknown functions. The members of *Rizhobium* regularly display considerable levels of genomic plasticity [64]. Hence, acquiring these genomics features in the WD29 strain may be linked with the symbiotic interaction with *Zea mays* roots.

The genome of Rhizobium is highly exposed to evolution by several mechanisms [65]. Here, genomic analysis evidenced that the WD29 strain harbors several insertion sequences (IS), but plasmids and phages were not detected. IS had a critical role in the appearance of accessory genes in our strain of study, as these elements are involved in events of gene duplication [66]. Our findings suggest that the random duplication promoted by IS is intensely engaged in the evolution of *R. miluonense* WD29. Moreover, *R. miluonense*, like its closely related species, exhibits predominant gene loss events. While genome evolution is frequently observed in obligate symbionts [67,68], and although *Rhizobium* species are not obligate symbionts, the plant-host interaction could be involved in the gene loss event observed. Nonetheless, the few gene gain events likely provide certain adaptive advantages related to plant-host interaction [69].

A particularly significant aspect is the presence of exopolysaccharide biosynthesis genes, which are crucial for biofilm formation and establishing stable root system associations [77-79]. These polysaccharides protect bacteria from adverse environmental conditions and enable root surface colonization. By synthesizing EPS, *R. miluonense* WD29 can establish a biofilm and adhere to the root surface, facilitating root system colonization and symbiotic relationship establishment. Furthermore, the EPS biosynthesis genes form a microaerophilic environment essential for nitrogen fixation [80-82]. Our results show a direct correlation between increased galactose concentration, EPS production, and nitrogen fixation rates, suggesting that EPS creates optimal conditions for nitrogenase activity.

The genomic analysis of *R. miluonense* WD29 highlights its predicted multifunctional roles in soil health and biocontrol. These predictions suggest ways of degrading persistent contaminants, remediating heavy metal contamination, fixing atmospheric nitrogen, and producing antimicrobial compounds to outcompete pathogenic bacteria. Additionally, the bacterium produces chitinase for fungal biocontrol and extracellular polysaccharides to enhance plant interactions. Some of these features have been demonstrated in vitro, where *R. miluonense* WD29 showed chitinase activity and produced substantial amounts of extracellular polysaccharides. This indicates its potential to inhibit fungal growth and facilitate beneficial plant-microbe interactions directly.

Further research is needed to validate these functions in natural environments and understand their implications for sustainable agriculture (Figure 6). Future research should focus on empirically testing these bioinformatics-based predictions to realize their full potential in sustainable agriculture. Moreover, exploring native maize resources as a source of highly interesting microorganisms underscores the significance of understanding the holobiont concept and its implications for agricultural sustainability and innovation.

**Figure 6.**
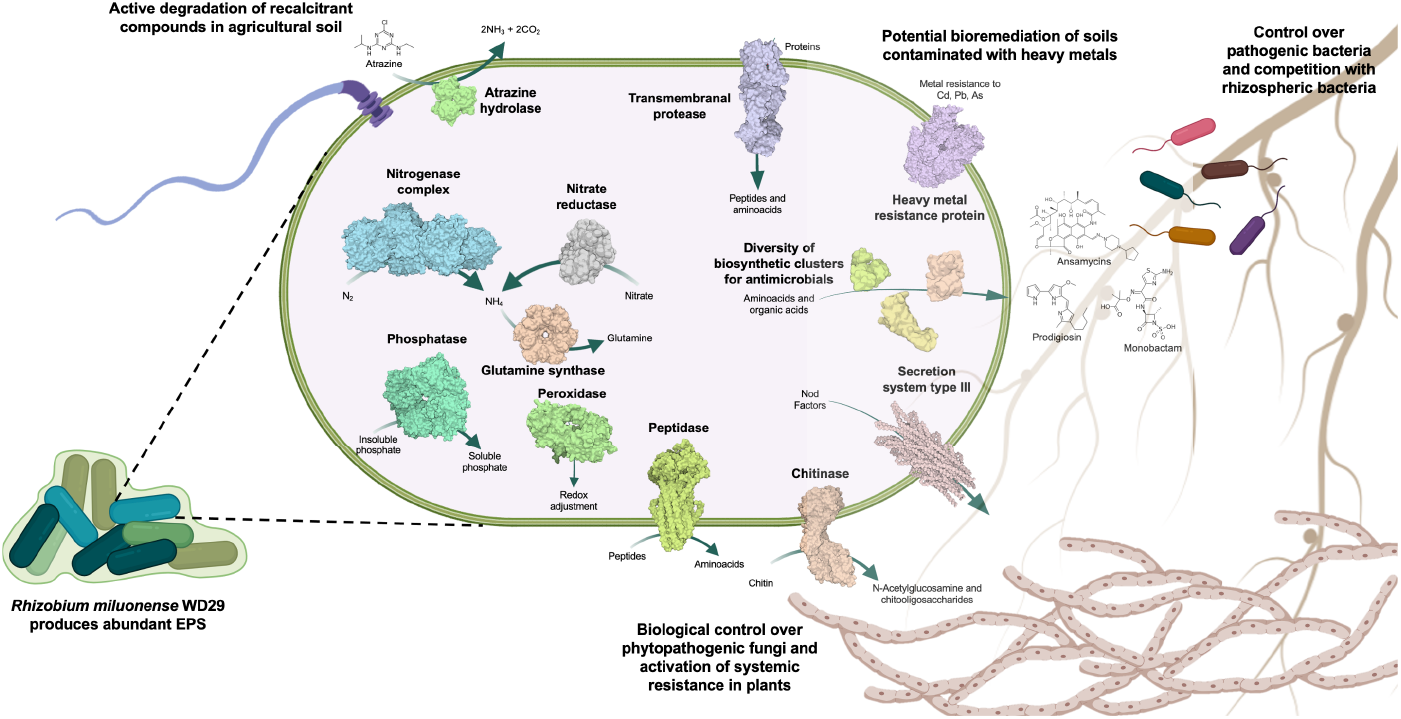
Multifunctional role of *Rhizobium miluonense* WD29 in soil health and biocontrol. Schematic representation of *Rhizobium miluonense* WD29’s roles in soil health and biocontrol. The bacterium contributes to the active degradation of recalcitrant compounds, such as atrazine, through specific enzymes like atrazine hydrolase. It aids in the bioremediation of heavy metal-contaminated soils by expressing heavy metal resistance proteins. *Rhizobium miluonense* WD29 produces various bioactive compounds, including antimicrobials, via diverse biosynthetic clusters and supports nitrogen fixation through a nitrogenase complex. It also secretes extracellular polysaccharides (EPS), deploys chitinase to control phytopathogenic fungi, and activates plant systemic resistance. It competes with pathogenic bacteria in the rhizosphere by secreting antimicrobial compounds. The scheme was performed in BioRender.

## 5. Conclusions

This study provides important insights into the genome of *R. miluonense* WD29, a bacterium with unique characteristics of free-living nitrogen fixation. The genomic and phenotypic analyses revealed distinctive features that enable efficient nitrogen fixation in free-living conditions and multiple plant growth-promoting traits. Identifying critical genes involved in exopolysaccharide biosynthesis and their correlation with nitrogen fixation efficiency represents a significant finding for understanding the molecular mechanisms of this process. The strain’s unique evolutionary adaptations suggest specialized mechanisms for plant-microbe interactions, particularly in non-leguminous associations. While field validation studies are still needed, these findings expand our understanding of rhizobial metabolic versatility and provide a foundation for developing sustainable agricultural practices. Future research should focus on validating these functions under field conditions and developing optimal application strategies for agricultural use. This research contributes to the growing body of knowledge about nitrogen-fixing bacteria and their potential role in enhancing crop productivity through environmentally friendly approaches.

## Supplementary Materials

### Author Contributions

EDVC and JOA performed the bacterial characterization and conducted the in vitro experiments; EDVC, JOA, and JAHG analyzed the bacterial genome; LVT and CHHR designed and coordinated the study; EDVC and CHHR wrote the first draft of the manuscript. All authors read, corrected, and approved the final manuscript.

### Funding

This research was funded by the Secretaría de Investigación y Posgrado-IPN (SIP 20220795, 20231480, and 20240945).

### Data Availability Statement

The *Rhizobium miluonense* WD29 genome sequence presented herein was deposited in GenBank under BioProject (XXXX) and BioSample (XXXX).

## Acknowledgments

All authors thank the review service for the English version that Sofía Marteli Mulia Castro conducted. EDVC is grateful to the Consejo Nacional de Humanidades Ciencia y Tecnología (CONAHCyT) and BEIFI for students. JAHG, LVT, and CHHR are fellows of EDI-IPN, COFFA-IPN, and SNI-CONAHCyT.

## Disclaimer/Publisher’s Note

The statements, opinions and data contained in all publications are solely those of the individual author(s) and contributor(s) and not of MDPI and/or the editor(s). MDPI and/or the editor(s) disclaim responsibility for any injury to people or property resulting from any ideas, methods, instructions or products referred to in the content.

## References

1. Ohyama, T. Nitrogen as a Major Essential Element of Plants. Nitrogen Assim. Plants 2010, 37, 1–17.

2. Mills, H.A.; Jones, J.B. Nutrient Deficiencies and Toxicities in Plants: Nitrogen. J. Plant Nutr. 1979, 1, 101–122. 10.1080/01904167909362704

3. Grotz, N.; Guerinot, M.L. Limiting Nutrients: An Old Problem with New Solutions? Curr. Opin. Plant Biol. 2002, 5, 158–163. 10.1016/S1369-5266(02)00247-9

4. Zheng, Z.L. Carbon and Nitrogen Nutrient Balance Signaling in Plants. Plant Signal. Behav. 2009, 4, 584–591. 10.4161/psb.4.7.8540

5. Cocking, E.C. Endophytic Colonization of Plant Roots by Nitrogen-Fixing Bacteria. Plant Soil 2003, 252, 169– 175. 10.1023/A:1024106605806

6. Bhattacharjee, R.B.; Singh, A.; Mukhopadhyay, S.N. Use of Nitrogen-Fixing Bacteria as Biofertiliser for Non-Legumes: Prospects and Challenges. Appl. Microbiol. Biotechnol. 2008, 80, 199–209. 10.1007/s00253-008-1567-2

7. Franche, C.; Lindström, K.; Elmerich, C. Nitrogen-Fixing Bacteria Associated with Leguminous and Non-Leguminous Plants. Plant Soil 2009, 321, 35–59. 10.1007/s11104-008-9833-8

8. Lindström, K.; Mousavi, S.A. Effectiveness of Nitrogen Fixation in Rhizobia. Microb. Biotechnol. 2020, 13, 1314– 1335. 10.1111/1751-7915.13517

9. Martínez-Espinosa, R.M.; Cole, J.A.; Richardson, D.J.; Watmough, N.J. Enzymology and Ecology of the Nitrogen Cycle. Biochem. Soc. Trans. 2011, 39, 175–178. 10.1042/BST0390175

10. Masson-Boivin, C.; Giraud, E.; Perret, X.; Batut, J. Establishing Nitrogen-Fixing Symbiosis with Legumes: How Many Rhizobium Recipes? Trends Microbiol. 2009, 17, 458–466. 10.1016/j.tim.2009.07.004

11. Geurts, R.; Bisseling, T. Rhizobium Nod Factor Perception and Signalling. Plant Cell 2002, 14, S239–S249. 10.1105/tpc.002451

12. Pereg, L.; Morugán-Coronado, A.; McMillan, M.; García-Orenes, F. Restoration of Nitrogen Cycling Community in Grapevine Soil by a Decade of Organic Fertilization. Soil Tillage Res. 2018, 179, 11–19. 10.1016/j.still.2018.01.007

13. Gómez-Godínez, L.J.; Fernandez-Valverde, S.L.; Martinez Romero, J.C.; Martínez-Romero, E. Metatranscriptomics and Nitrogen Fixation from the Rhizoplane of Maize Plantlets Inoculated with a Group of PGPRs. Syst. Appl. Microbiol. 2019, 42, 517–525. 10.1016/j.syapm.2019.05.003

14. Qin, S.; Pang, Y.; Clough, T.; Wrage-Mönnig, N.; Hu, C.; Zhang, Y.; Zhou, S.; Fang, Y. N2 Production via Aerobic Pathways May Play a Significant Role in Nitrogen Cycling in Upland Soils. Soil Biol. Biochem. 2017, 108, 36–40. 10.1016/j.soilbio.2017.01.019

15. Thamer, S.; Schädler, M.; Bonte, D.; Ballhorn, D.J. Dual Benefit from a Belowground Symbiosis: Nitrogen Fixing Rhizobia Promote Growth and Defense against a Specialist Herbivore in a Cyanogenic Plant. Plant Soil 2011, 341, 209–219. 10.1007/s11104-010-0635-4

16. Pérez-Montaño, F.; Jiménez-Guerrero, I.; Acosta-Jurado, S.; Navarro-Gómez, P.; Ollero, F.J.; Ruiz-Sainz, J.E.; López-Baena, F.J.; Vinardell, J.M. A Transcriptomic Analysis of the Effect of Genistein on Sinorhizobium fredii HH103 Reveals Novel Rhizobial Genes Putatively Involved in Symbiosis. Sci. Rep. 2016, 6, 31592. 10.1038/srep31592

17. Sachs, J.L.; Quides, K.W.; Wendlandt, C.E. Legumes versus Rhizobia: A Model for Ongoing Conflict in Symbiosis. New Phytol. 2018, 219, 1199–1206. 10.1111/nph.15222

18. Long, S.R. Rhizobium-Legume Nodulation: Life Together in the Underground. Cell 1989, 56, 203–214. 10.1016/0092-8674(89)90893-3

19. Kim, J.; Rees, D.C. Nitrogenase and Biological Nitrogen Fixation. Biochemistry 1994, 33, 389–397. 10.1021/bi00168a001

20. Ludwig, R.A. Rhizobium Free-Living Nitrogen Fixation Occurs in Specialized Nongrowing Cells. Proc. Natl. Acad. Sci. USA 1984, 81, 1566–1569. 10.1073/pnas.81.5.1566

21. Dunican, L.K.; Tierney, A.B. Genetic Transfer of Nitrogen Fixation from Rhizobium trifolii to Klebsiella aerogenes. Biochem. Biophys. Res. Commun. 1974, 57, 62–72. 10.1016/S0006-291X(74)80357-8

22. Domenzain, C.; Camarena, L.; Osorio, A.; Dreyfus, G.; Poggio, S. Evolutionary Origin of the Rhodobacter sphaeroides Specialized RpoN Sigma Factors. FEMS Microbiol. Lett. 2012, 327, 93–102. 10.1111/j.1574-6968.2011.02459.x.

23. Yuan, K.; Reckling, M.; Ramirez, M.D.A.; Djedidi, S.; Fukuhara, I.; Ohyama, T.; Yokoyama, T.; Bellingrath-Kimura, S.D.; Halwani, M.; Egamberdieva, D. Characterization of Rhizobia for the Improvement of Soybean Cultivation at Cold Conditions in Central Europe. Microb. Environ. 2020, 35, ME19124. 10.1264/jsme2.ME19124

24. Getahun, A.; Muleta, D.; Assefa, F.; Kiros, S. Field Application of Rhizobial Inoculants in Enhancing Faba Bean Production in Acidic Soils: An Innovative Strategy to Improve Crop Productivity. In Salt Stress, Microbes, and Plant Interactions: Causes and Solution; Akhtar, M.S., Ed.; Springer: Singapore, 2019; pp. 147–180.

25. Gu, C.T.; Wang, E.T.; Tian, C.F.; Han, T.X.; Chen, W.F.; Sui, X.H.; Chen, W.X. Rhizobium miluonense Sp. Nov., a Symbiotic Bacterium Isolated from Lespedeza Root Nodules. Int. J. Syst. Evol. Microbiol. 2008, 58, 1364–1368. 10.1099/ijs.0.65661-0

26. Rocha, S.M.B.; Do Amorim, M.R.; Costa, M.K.L.; Da Silva Saraiva, T.C.; Costa, R.M.; Antunes, J.E.L.; De Souza Oliveira, L.M.; De Alcantara Neto, F.; De Medeiros, E.V.; De Araujo Pereira, A.P. Tolerance and Reduction of Chromium by Bacterial Strains. Arch. Microbiol. 2022, 204, 730. 10.1007/s00203-022-03329-3

27. Arpiwi, N.L.; Yan, G.; Barbour, E.L.; Plummer, J.A.; Watkin, E. Phenotypic and Genotypic Characterisation of Root Nodule Bacteria Nodulating Millettia pinnata (L.) Panigrahi, a Biodiesel Tree. Plant Soil 2013, 367, 363–377. 10.1007/s11104-012-1472-4

28. Rios-Galicia, B.; Villagómez-Garfias, C.; De La Vega-Camarillo, E.; Guerra-Camacho, J.E.; Medina-Jaritz, N.; Arteaga-Garibay, R.I.; Villa-Tanaca, L.; Hernández-Rodríguez, C. The Mexican Giant Maize of Jala Landrace Harbour Plant-Growth-Promoting Rhizospheric and Endophytic Bacteria. 3 Biotech 2021, 11, 447. 10.1007/s13205-021-02983-6

29. William, S.; Feil, H.; Copeland, A. Bacterial Genomic DNA Isolation Using CTAB. Sigma 2012, 50, 6876.

30. Andrews, S. FastQC: A Quality Control Tool for High Throughput Sequence Data. 2010.

31. Bolger, A.M.; Lohse, M.; Usadel, B. Trimmomatic: A Flexible Trimmer for Illumina Sequence Data. Bioinformatics 2014, 30, 2114–2120. 10.1093/bioinformatics/btu170

32. Bankevich, A.; Nurk, S.; Antipov, D.; Gurevich, A.A.; Dvorkin, M.; Kulikov, A.S.; Lesin, V.M.; Nikolenko, S.I.; Pham, S.; Prjibelski, A.D.; Pyshkin, A.V.; Sirotkin, A.V.; Vyahhi, N.; Tesler, G.; Alekseyev, M.A.; Pevzner, P.A. SPAdes: A New Genome Assembly Algorithm and Its Applications to Single-Cell Sequencing. J. Comput. Biol. 2012, 19, 455–477. 10.1089/cmb.2012.0021

33. Gurevich, A.; Saveliev, V.; Vyahhi, N.; Tesler, G. QUAST: Quality Assessment Tool for Genome Assemblies. Bioinformatics 2013, 29, 1072–1075. 10.1093/bioinformatics/btt086

34. Simão, F.A.; Waterhouse, R.M.; Ioannidis, P.; Kriventseva, E.V.; Zdobnov, E.M. BUSCO: Assessing Genome Assembly and Annotation Completeness with Single-Copy Orthologs. Bioinformatics 2015, 31, 3210–3212. 10.1093/bioinformatics/btv351

35. Aziz, R.K.; Bartels, D.; Best, A.A.; DeJongh, M.; Disz, T.; Edwards, R.A.; Formsma, K.; Gerdes, S.; Glass, E.M.; Kubal, M.; Meyer, F.; Olsen, G.J.; Olson, R.; Osterman, A.L.; Overbeek, R.A.; McNeil, L.K.; Paarmann, D.; Paczian, T.; Parrello, B.; Pusch, G.D.; et al. The RAST Server: Rapid Annotations Using Subsystems Technology. BMC Genom. 2008, 9, 75. 10.1186/1471-2164-9-75

36. Seemann, T. Prokka: Rapid Prokaryotic Genome Annotation. Bioinformatics 2014, 30, 2068–2069. 10.1093/bioinformatics/btu153

37. Tatusova, T.; DiCuccio, M.; Badretdin, A.; Chetvernin, V.; Nawrocki, E.P.; Zaslavsky, L.; Lomsadze, A.; Pruitt, K.D.; Borodovsky, M.; Ostell, J. NCBI Prokaryotic Genome Annotation Pipeline. Nucleic Acids Res. 2016, 44, 6614–6624. 10.1093/nar/gkw569

38. Blin, K.; Shaw, S.; Steinke, K.; Villebro, R.; Ziemert, N.; Lee, S.Y.; Medema, M.H.; Weber, T. antiSMASH 5.0: Updates to the Secondary Metabolite Genome Mining Pipeline. Nucleic Acids Res. 2019, 47, W81–W87. 10.1093/nar/gkz310

39. Skinnider, M.A.; Dejong, C.A.; Rees, P.N.; Johnston, C.W.; Li, H.; Webster, A.L.H.; Wyatt, M.A.; Magarvey, N.A. Genomes to Natural Products PRediction Informatics for Secondary Metabolomes (PRISM). Nucleic Acids Res. 2015, 43, 9645–9662. 10.1093/nar/gkv1012

40. Goris, J.; Konstantinidis, K.T.; Klappenbach, J.A.; Coenye, T.; Vandamme, P.; Tiedje, J.M. DNA–DNA Hybridization Values and Their Relationship to Whole-Genome Sequence Similarities. Int. J. Syst. Evol. Microbiol. 2007, 57, 81–91. 10.1099/ijs.0.64483-0

41. Pritchard, L.; Cock, P.; Esen, Ö. PyANI: Python Module for Average Nucleotide Identity Analysis. Version 0.2.8. 2019.

42. Avram, O.; Rapoport, D.; Portugez, S.; Pupko, T. M1CR0B1AL1Z3R—A User-Friendly Web Server for the Analysis of Large-Scale Microbial Genomics Data. Access Microbiol. 2020, 2, po1014. 10.1099/acmi.ac2020.po1014

43. Chen, C.; Chen, H.; Zhang, Y.; Thomas, H.R.; Frank, M.H.; He, Y.; Xia, R. TBtools: An Integrative Toolkit Developed for Interactive Analyses of Big Biological Data. Mol. Plant 2020, 13, 1194–1202. 10.1016/j.molp.2020.06.009

44. Sun, J.; Lu, F.; Luo, Y.; Bie, L.; Xu, L.; Wang, Y. OrthoVenn3: An Integrated Platform for Exploring and Visualizing Orthologous Data across Genomes. Nucleic Acids Res. 2023, 51, W397–W403. 10.1093/nar/gkad313

45. Darling, A.C.E.; Mau, B.; Blattner, F.R.; Perna, N.T. Mauve: Multiple Alignment of Conserved Genomic Sequence with Rearrangements. Genome Res. 2004, 14, 1394–1403. 10.1101/gr.2289704

46. Rahimlou, S.; Bahram, M.; Tedersoo, L. Phylogenomics Reveals the Evolution of Root Nodulating Alpha- and Beta-Proteobacteria (Rhizobia). Microbiol. Res. 2021, 250, 126788. 10.1016/j.micres.2021.126788

47. Afzal, I.; Iqrar, I.; Shinwari, Z.K.; Yasmin, A. Plant Growth-Promoting Potential of Endophytic Bacteria Isolated from Roots of Wild Dodonaea viscosa L. Plant Growth Regul. 2017, 81, 399–408. 10.1007/s10725-016-0216-5

48. Hardy, R.W.F.; Holsten, R.D.; Jackson, E.K.; Burns, R.C. The Acetylene-Ethylene Assay for N2 Fixation: Laboratory and Field Evaluation. Plant Physiol. 1968, 43, 1185–1207. 10.1104/pp.43.8.1185

49. Kanehisa, M.; Araki, M.; Goto, S.; Hattori, M.; Hirakawa, M.; Itoh, M.; Katayama, T.; Kawashima, S.; Okuda, S.; Tokimatsu, T.; et al. KEGG for Linking Genomes to Life and the Environment. Nucleic Acids Res. 2007, 36, D480– D484. 10.1093/nar/gkm882

50. Kanehisa, M.; Sato, Y.; Morishima, K. BlastKOALA and GhostKOALA: KEGG Tools for Functional Characterization of Genome and Metagenome Sequences. J. Mol. Biol. 2016, 428, 726–731. 10.1016/j.jmb.2015.11.006

51. Jones, P.; Binns, D.; Chang, H.Y.; Fraser, M.; Li, W.; McAnulla, C.; McWilliam, H.; Maslen, J.; Mitchell, A.; Nuka, G.; et al. InterProScan 5: Genome-Scale Protein Function Classification. Bioinformatics 2014, 30, 1236–1240. 10.1093/bioinformatics/btu031

52. Sievers, F.; Higgins, D.G. Clustal Omega. Curr. Protoc. Bioinformatics 2014, 48, 3.13.1–3.13.16. 10.1002/0471250953.bi0313s48

53. Procter, J.B.; Carstairs, G.M.; Soares, B.; Mourão, K.; Ofoegbu, T.C.; Barton, D.; Lui, L.; Menard, A.; Sherstnev, N.; Roldan-Martinez, D.; et al. Alignment of Biological Sequences with Jalview. In Multiple Sequence Alignment; Katoh, K., Ed.; Methods in Molecular Biology; Springer: New York, NY, USA, 2021; Volume 2231, pp. C1–C1.

54. Skolnick, J.; Gao, M.; Zhou, H.; Singh, S. AlphaFold 2: Why It Works and Its Implications for Understanding the Relationships of Protein Sequence, Structure, and Function. J. Chem. Inf. Model. 2021, 61, 4827–4831. 10.1021/acs.jcim.1c01114

55. Mahram, A.; Herbordt, M.C. NCBI BLASTP on High-Performance Reconfigurable Computing Systems. ACM Trans. Reconfigurable Technol. Syst. 2015, 7, 1–20. 10.1145/2629691

56. Remigi, P.; Zhu, J.; Young, J.P.W.; Masson-Boivin, C. Symbiosis within Symbiosis: Evolving Nitrogen-Fixing Legume Symbionts. Trends Microbiol. 2016, 24, 63–75. 10.1016/j.tim.2015.10.007

57. Chen, W.F.; Wang, E.T.; Ji, Z.J.; Zhang, J.J. Recent Development and New Insight of Diversification and Symbiosis Specificity of Legume Rhizobia: Mechanism and Application. J. Appl. Microbiol. 2021, 131, 553–563. 10.1111/jam.14960

58. Yang, J.; Lan, L.; Jin, Y.; Yu, N.; Wang, D.; Wang, E. Mechanisms Underlying Legume–Rhizobium Symbioses. J. Integr. Plant Biol. 2022, 64, 244–267. 10.1111/jipb.13207

59. Andrews, M.; Andrews, M.E. Specificity in Legume-Rhizobia Symbioses. Int. J. Mol. Sci. 2017, 18, 705. 10.3390/ijms18040705

60. García-Fraile, P.; Carro, L.; Robledo, M.; Ramírez-Bahena, M.H.; Flores-Félix, J.D.; Fernández, M.T.; Mateos, P.F.; Rivas, R.; Igual, J.M.; Martínez-Molina, E.; et al. Rhizobium Promotes Non-Legumes Growth and Quality in Several Production Steps: Towards a New Biofertilizer. Microb. Cell Fact. 2012, 11, 125. 10.1186/1475-2859-11-125

61. De Oliveira-Francesquini, J.P.; Hungria, M.; Savi, D.C.; Glienke, C.; Aluizio, R.; Kava, V.; Galli-Terasawa, L.V. Differential Colonization by Bioprospected Rhizobial Bacteria Associated with Common Bean in Different Cropping Systems. Can. J. Microbiol. 2017, 63, 682–689. 10.1139/cjm-2016-0784

62. Wisniewski-Dyé, F.; Drogue, B.; Borland, S.; Prigent-Combaret, C. Azospirillum-Plant Interaction: From Root Colonization to Plant Growth Promotion. Beneficial Plant-Microbial Interactions 2013, 237–269. 10.1201/b15251-11

63. Taktek, S.; St-Arnaud, M.; Piché, Y.; Fortin, J.A.; Antoun, H. Igneous Phosphate Rock Solubilization by Biofilm-Forming Mycorrhizobacteria and Hyphobacteria Associated with Rhizoglomus irregulare DAOM 197198. Mycorrhiza 2017, 27, 13–22. 10.1007/s00572-016-0726-z

64. Rosselli, R.; La Porta, N.; Muresu, R.; Stevanato, P.; Concheri, G.; Squartini, A. Pangenomics of the Symbiotic Rhizobiales: Core and Accessory Functions Across a Group Endowed with High Levels of Genomic Plasticity. Microorganisms 2021, 9, 407. 10.3390/microorganisms9020407

65. Epstein, B.; Tiffin, P. Comparative Genomics Reveals High Rates of Horizontal Transfer and Strong Purifying Selection on Rhizobial Symbiosis Genes. Proc. R. Soc. B 2021, 288, 20201804. 10.1098/rspb.2020.1804

66. Arashida, H.; Odake, H.; Sugawara, M.; Noda, R.; Kakizaki, K.; Ohkubo, S.; Mitsui, H.; Sato, S.; Minamisawa, K. Evolution of Rhizobial Symbiosis Islands through Insertion Sequence-Mediated Deletion and Duplication. ISME J. 2022, 16, 112–121. 10.1038/s41396-021-01035-4

67. McCutcheon, J.P.; Moran, N.A. Extreme Genome Reduction in Symbiotic Bacteria. Nat. Rev. Microbiol. 2012, 10, 13–26. 10.1038/nrmicro2670

68. Zhao, R.; Liu, L.X.; Zhang, Y.Z.; Jiao, J.; Cui, W.J.; Zhang, B.; Wang, X.L.; Li, M.L.; Chen, Y.; Xiong, Z.Q.; et al. Adaptive Evolution of Rhizobial Symbiotic Compatibility Mediated by Co-Evolved Insertion Sequences. ISME J. 2018, 12, 101–111. 10.1038/ismej.2017.136

69. Mower, J.P.; Stefanović, S.; Young, G.J.; Palmer, J.D. Gene Transfer from Parasitic to Host Plants. Nature 2004, 432, 165–166. 10.1038/432165b

70. González, V.; Santamaría, R.I.; Bustos, P.; Hernández-González, I.; Medrano-Soto, A.; Moreno-Hagelsieb, G.; Janga, S.C.; Ramírez, M.A.; Jiménez-Jacinto, V.; Collado-Vides, J.; et al. The Partitioned Rhizobium etli Genome: Genetic and Metabolic Redundancy in Seven Interacting Replicons. Proc. Natl. Acad. Sci. USA 2006, 103, 3834– 3839. 10.1073/pnas.0508502103

71. Adato, O.; Ninyo, N.; Gophna, U.; Snir, S. Detecting Horizontal Gene Transfer between Closely Related Taxa. PLoS Comput. Biol. 2015, 11, e1004408. 10.1371/journal.pcbi.1004408

72. Kuan, K.B.; Othman, R.; Abdul Rahim, K.; Shamsuddin, Z.H. Plant Growth-Promoting Rhizobacteria Inoculation to Enhance Vegetative Growth, Nitrogen Fixation and Nitrogen Remobilisation of Maize under Greenhouse Conditions. PLoS ONE 2016, 11, e0152478. 10.1371/journal.pone.0152478

73. Castellano-Hinojosa, A.; Correa-Galeote, D.; Palau, J.; Bedmar, E.J. Isolation of N2-fixing Rhizobacteria from Lolium perenne and Evaluating their Plant Growth Promoting Traits. Biology 2016, 5, 25. 10.3390/biology5020025

74. Renoud, S.; Bouffaud, M.L.; Dubost, A.; Prigent-Combaret, C.; Legendre, L.; Moënne-Loccoz, Y.; Muller, D. Co-Occurrence of Rhizobacteria with Nitrogen Fixation and/or 1-Aminocyclopropane-1-Carboxylate Deamination Abilities in the Maize Rhizosphere. FEMS Microbiol. Ecol. 2020, 96, fiaa062. 10.1093/femsec/fiaa062

75. Gastélum, G.; Aguirre-von-Wobeser, E.; De La Torre, M.; Rocha, J. Interaction Networks Reveal Highly Antagonistic Endophytic Bacteria in Native Maize Seeds from Traditional milpa Agroecosystems. Environ. Microbiol. 2022, 24, 5583–5595. 10.1111/1462-2920.16189

76. Breedt, G.; Labuschagne, N.; Coutinho, T.A. Seed Treatment with Selected Plant Growth-promoting Rhizobacteria Increases Maize Yield in the Field. Ann. Appl. Biol. 2017, 171, 229–236. 10.1111/aab.12366

77. Zhang, N.; Yang, D.; Wang, D.; Miao, Y.; Shao, J.; Zhou, X.; Xu, Z.; Li, Q.; Feng, H.; Li, S.; et al. Whole Transcriptomic Analysis of the Plant-Beneficial Rhizobacterium Bacillus amyloliquefaciens SQR9 during Enhanced Biofilm Formation Regulated by Maize Root Exudates. BMC Genomics 2015, 16, 685. 10.1186/s12864-015-1825-5

78. Pérez-Montaño, F.; Jiménez-Guerrero, I.; Del Cerro, P.; Baena-Ropero, I.; López-Baena, F.J.; Ollero, F.J.; Bellogín, R.; Lloret, J.; Espuny, R. The Symbiotic Biofilm of Sinorhizobium fredii SMH12, Necessary for Successful Colonization and Symbiosis of Glycine max cv Osumi, Is Regulated by Quorum Sensing Systems and Inducing Flavonoids via NodD1. PLoS ONE 2014, 9, e105901. 10.1371/journal.pone.0105901

79. Bonkowski, M.; Tarkka, M.; Razavi, B.S.; Schmidt, H.; Blagodatskaya, E.; Koller, R.; Yu, P.; Knief, C.; Hochholdinger, F.; Vetterlein, D. Spatiotemporal Dynamics of Maize (Zea mays L.) Root Growth and Its Potential Consequences for the Assembly of the Rhizosphere Microbiota. Front. Microbiol. 2021, 12, 619499. 10.3389/fmicb.2021.619499

80. Meneses, C.H.S.G.; Rouws, L.F.M.; Simões-Araújo, J.L.; Vidal, M.S.; Baldani, J.I. Exopolysaccharide Production Is Required for Biofilm Formation and Plant Colonization by the Nitrogen-Fixing Endophyte Gluconacetobacter diazotrophicus. MPMI 2011, 24, 1448–1458. 10.1094/MPMI-05-11-0127

81. Wang, D.; Xu, A.; Elmerich, C.; Ma, L.Z. Biofilm Formation Enables Free-Living Nitrogen-Fixing Rhizobacteria to Fix Nitrogen under Aerobic Conditions. ISME J. 2017, 11, 1602–1613. 10.1038/ismej.2017.30

82. He, X.; Li, Q.; Wang, N.; Chen, S. Effects of an EPS Biosynthesis Gene Cluster of Paenibacillus polymyxa WLY78 on Biofilm Formation and Nitrogen Fixation under Aerobic Conditions. Microorganisms 2021, 9, 289. 10.3390/microorganisms9020289

